# UV-induced translation capacity is regulated developmentally in fungi

**DOI:** 10.64898/2026.08.04.742702

**Authors:** Quyen Hoang, Shira Milo, Dibya Mukherjee, Shay Covo

**Author notes:** Corresponding author Shay Covo. **Email:**.

## Abstract

In fast-growing cells, most RNA molecules encode ribosomal subunits. During growth arrest caused by DNA damage, ribosome biogenesis is reduced because it is a highly energy-demanding process. Here, we report that, in two filamentous fungi, UV radiation significantly induces ribosome levels and translation capacity. This UV-induced response is developmentally regulated and occurs after filament formation. Translation induction modestly but significantly enhances both UV survival and repair. While some of the mRNAs that are differentially associated with ribosomes following UV exposure are associated with DNA repair, most of them belong to modules involved in protein folding and gene expression. TOR signaling is required for the immediate UV induction of ribosome biogenesis genes. However, we provide evidence that UV induces mechanisms that operate either upstream of or in parallel with TOR. These findings suggest a novel type of response to DNA damage, in which the primary aim is to protect the integrity of gene expression from the detrimental effects of UV radiation on transcription.

## Introduction

Ultraviolet (UV) light is an important determinant in the ecology and physiology of both prokaryotic and eukaryotic microbes (Albarracín, Pathak et al. 2012, Palmer, Drees et al. 2018). While UV can damage several biochemical components, probably the most sensitive is DNA, since mutants in DNA repair are extremely sensitive to UV (Cleaver 2016). The most common UV lesions are cyclobutane pyrimidine dimers (CPDs) and 6–4 UV photoproducts, both of which significantly distort the DNA helix (Rastogi, Richa et al. 2010). This severe disruption of DNA structure interrupts DNA replication, leading to stalled DNA replication forks and cell death. UV is also mutagenic; primarily, it causes point mutations, most commonly C-to-T transitions (Laughery, Brown et al. 2020). In addition to its negative effect on DNA replication, UV inhibits transcription (Nieto Moreno, Olthof et al. 2023).

Several DNA repair mechanisms suppress the genotoxic and mutagenic effects of UV. These mechanisms can be divided into those that remove lesions from the entire genome and those that remove lesions from transcribed genes. UV damage repair mechanisms in ascomycete fungi include UV damage endonuclease (UVDE) and photolyase, which both specifically recognize UV lesions and act globally across the entire genome (McCready, Osman et al. 2000, Goldman, McGuire et al. 2002, Inoue 2011). Nucleotide excision repair (NER) recognizes DNA helix distortions by scanning the genome using the proteins Ddb1 and Xpc (global NER) (Lindahl, Karran et al. 1997, Moser, Volker et al. 2005, Scrima, Fischer et al. 2011). Alternatively, the lesions are recognized by stalled RNA polymerase II with the assistance of proteins Csa and Csb, a process known as transcription-coupled (TCR) NER (Lindahl, Karran et al. 1997, Lindahl and Wood 1999, Sancar and Reardon 2004, Moser, Volker et al. 2005). TCR facilitates the resumption of transcription following UV exposure more rapidly than if only global DNA repair mechanisms were operating. UV-induced transcription stalling is probably a severe challenge, since eukaryotic cells evolved a last-resort solution to this problem by degrading stalled RNA polymerase II. In fact, TCR is also required for the release of stalled RNA polymerase I (Daniel, Cerutti et al. 2018, Nieto Moreno, Olthof et al. 2023).

The bacterial SOS response is the model of the cellular response to UV (Maslowska, Makiela-Dzbenska et al. 2019). During SOS activation, there is an orchestrated induction of UV repair and mutagenesis genes. The SOS response induces, at the mRNA level, the expression of DNA repair, recombination, and DNA damage tolerance genes, some of which increase up to 100-fold (Maslowska, Makiela-Dzbenska et al. 2019). A comprehensive, SOS-like induction of DNA repair genes has never been demonstrated in fungi. Nevertheless, some DNA repair/DNA damage tolerance genes are induced by UV in *Saccharomyces cerevisiae*, *Schizosaccharomyces pombe*, and *Neurospora crassa* (Davey, Nass et al. 1997, Sakuraba, Schroeder et al. 2000, Fu, Pastushok et al. 2008).

Several attempts were made to characterize the UV response in various eukaryotic organisms under different exposure and recovery conditions. In general, the transcriptional response was variable and complex, highlighting responses to reactive oxygen species, flavonoid biosynthesis, and cell cycle control (Wade, Poorey et al. 2009, Pontin, Piccoli et al. 2010, Guo, Yu et al. 2019, Milo-Cochavi, Adar et al. 2019, Milo, Namawejje et al. 2024). A major part of the eukaryotic response to UV and other forms of DNA damage is cell cycle arrest, which allows cell division to occur only when genome integrity is not severely compromised (Carr 2002, Sertic, Pizzi et al. 2012). Thus, segregation of partly replicated DNA into the daughter cells is avoided.

In many aspects, the cellular response to DNA damage is a particular case of stress response that leads to growth arrest. In fast-growing yeast cells, up to 60% of genes are involved in ribosome biogenesis; however, this process is resource-intensive. Therefore, under-expression of ribosomal proteins is a sign of growth arrest (Warner 1999).

Translation is reduced in response to UV and oxidative stress, which also causes DNA damage (Ying and Khaperskyy 2020, Picazo and Molin 2021, Meydan, Barros et al. 2023). The effect of DNA damage on ribosome biogenesis was demonstrated at the cellular and molecular levels. In yeast, Xbp1, which suppresses the expression of ribosomal proteins and ribosome biogenesis, is activated by the DNA damage response (Tkach, Yimit et al. 2012, Miles, Li et al. 2013). In addition, UV-induced DNA lesions specifically cause dissociation of yeast RNA polymerases I (Tremblay, Charton et al. 2014). Further evidence supports a link between DNA repair and ribosome biosynthesis (Ogawa and Baserga 2017). In humans, under normal growth conditions, several DNA repair proteins, such as the Ape1 endonuclease and the Werner syndrome helicase, are localized to the nucleolus. They are thought to be involved in ribosome biogenesis. Once the DNA damage response is triggered, these repair proteins dissociate from the nucleolus (Lirussi, Antoniali et al. 2012). The opposite is also true: some ribosome biogenesis genes that leave the nucleolus following DNA damage participate in DNA repair (Ogawa and Baserga 2017).

Here we present our study on the UV response of two filamentous ascomycete fungi: *Fusarium mangiferae* and *F. oxysporum*, both of which are plant pathogens (Marasas, Ploetz et al. 2006, Michielse and Rep 2009, Ploetz 2015). The life cycle of these organisms starts with the germination of a single cell, leading to development from a single-nucleus spore (conidium) to a filamentous hypha. With time, the hypha elongates and branches to form a developed mycelium (Li, He et al. 2025). Only a few of the nuclei within the mycelium are mitotically active; the rest are dormant (Ruiz-Roldán, Köhli et al. 2010). Based on the above differences between fungal spores and filaments, we thought to study the response to UV at different stages of fungal development.

We previously reported the analysis of the transcriptomic response of *F. mangiferae* and *F. oxysporum* to UV under two developmental stages: 8 and 14 hours post-inoculation (hpi), while in the former stage, germination occurs with only one or two nuclei; in the latter stage, the filament is established with 10 or more nuclei (Milo-Cochavi, Adar et al. 2019, Milo, Namawejje et al. 2024). As expected, the expression of genes related to translation was downregulated following UV irradiation of fungi at 8 hpi. Surprisingly, we observed a remarkable induction of translation-related genes in both fungi when they were irradiated at 14 hpi (Milo-Cochavi, Adar et al. 2019, Milo, Namawejje et al. 2024).

There are two nutrient-sensing pathways that induce ribosome biogenesis: the target of rapamycin (TOR) and the cAMP-activated kinase PKA (Guerra, Vuillemenot et al. 2022, Plank 2022). There is considerable crosstalk and redundancy between the pathways, as well as some specificity. Both pathways activate, among others, Sfp1 and Stb3, which drive ribosome biogenesis, and deactivate Tod6 and Dot6, which suppresses ribosome biogenesis (Plank 2022).

Here, we show that ribosome abundance is markedly increased in response to UV and that translation capacity is markedly induced in both fungi. We found that UV-induced translation capacity is important for both UV repair and survival. Our research proposes a novel strategy for responding to UV lesions, about five decades after SOS was identified in bacteria.

## Results

### Transcriptome analysis of *Fusarium mangiferae* irradiated with UV 14 hpi reveals induction of ribosome biogenesis genes

Using network analysis, we reanalyzed the RNA-seq data for *F. mangiferae* irradiated with UV at 14 hours post-inoculation (hpi), grouping several recovery times (Milo-Cochavi, Adar et al. 2019, Milo, Namawejje et al. 2024). Among the biological modules that were most significantly induced by UV at 14 hpi were translation and ribosome biogenesis (Fig. S1). This observation is surprising, since exposure to DNA damage in Fusarium is usually associated with reduced ribosome biogenesis due to its negative effect on cell growth (Tremblay, Charton et al. 2014, Milo-Cochavi, Adar et al. 2019, Milo-Cochavi, Pareek et al. 2019). Therefore, we first confirmed that UV inflicts growth delay under these conditions. Without irradiation, the number of nuclei in the filament increased steadily every two hours throughout the experiment. When filaments were irradiated at 14 hpi, *F. mangiferae* showed no significant change in the number of nuclei during the first 4 hours. *Fusarium oxysoirum* f. sp. *lycopersici* (*Fol*) exhibited no change during the first 2 hours, followed by a modest increase after 4 hours (Fig. S2).

In addition to ribosome biogenesis genes, we found that pyrimidine biosynthesis and mitochondrion-related processes were both upregulated at 200 J/m^2^ exposure. Modules related to protein degradation were downregulated when cells were exposed to 200 J/m^2^ (Fig. S1). The increase in expression of pyrimidine biosynthesis, mitochondrial-related, and ribosome biogenesis genes following UV exposure suggests that rNTPs are consumed for substantial ribosome assembly.

### UV induces ribosomes in *Fusarium* species irradiated in a developmentally regulated manner

To test whether there is a ribosome buildup following irradiation, we irradiated 14 hpi filaments and collected samples 2 hours after irradiation. Extracts of the samples were separated on a sucrose gradient, and ribosomal RNA band intensities for each fraction obtained were measured (see Materials and Methods). We found that following irradiation, the amount of ribosomes in *F. mangiferae* increased (Fig. 1A), even at a low UV dose (50 J/m^2^). Specifically, we observed an increase in the RNA bands corresponding to ribosomes in the bottom fractions of the gradient, indicating an increase in the amount of polysomes.

**Figure 1.**
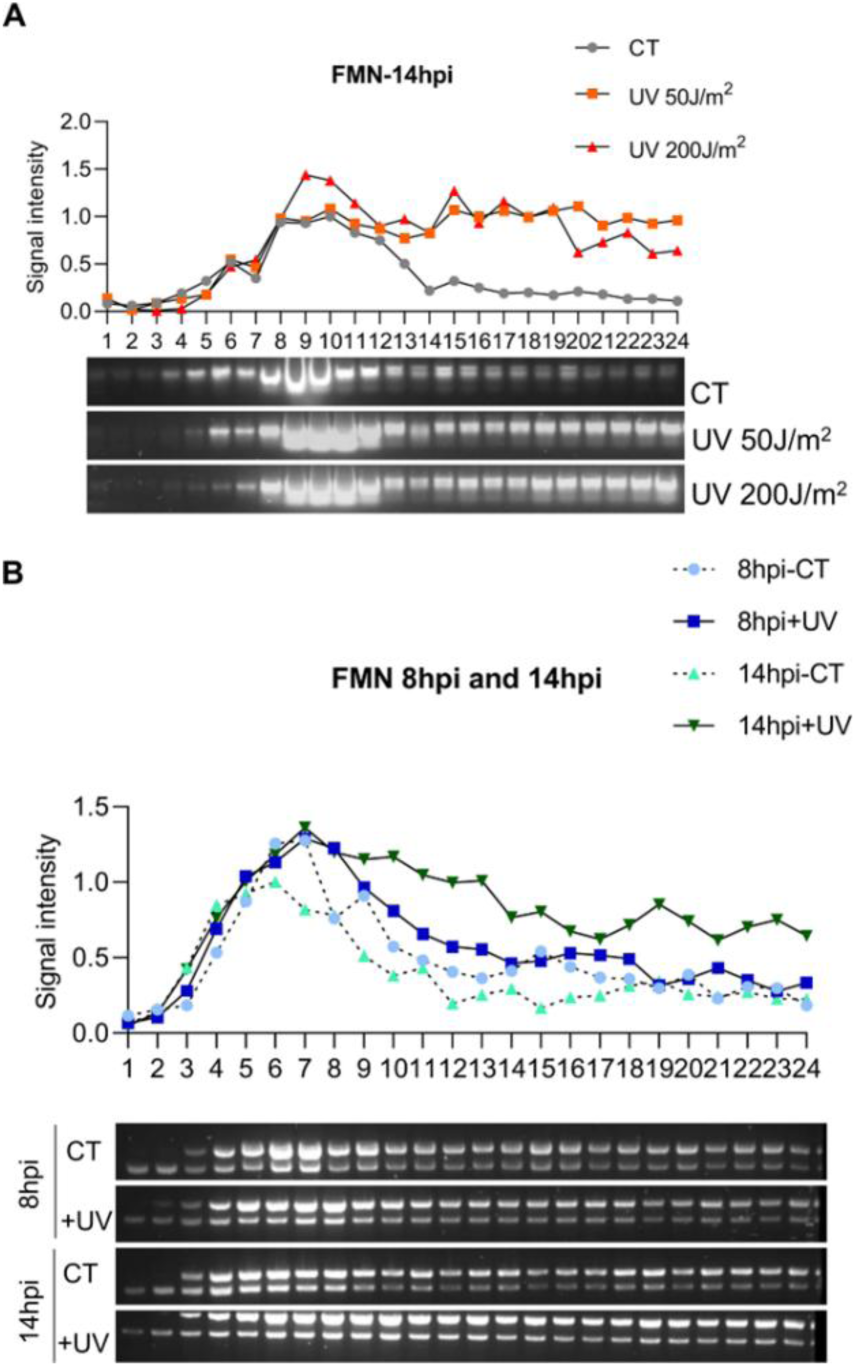
UV induces ribosome biogenesis in a developmentally dependent manner in *F. mangiferae*. Conidia of *F. mangiferae* (FMN) were inoculated for 8 (germlings) or 14 h (filaments) in PDB medium. Fungal filaments (A) or fungal germlings (B) were irradiated with UV or not (CT). The fungi were recovered for two hours in water. After recovery, the cultures were lysed and loaded on a sucrose cushion, followed by separation on a gradient as described under Methods. The RNA samples from each fraction (X axis) of the sucrose gradients were separated on 2% agarose gels, and the ribosome intensity was calculated based on the strongest band signal in the control sample (signal intensity as 1).

We then asked whether this UV induction of ribosome biosynthesis depends on the developmental stage of the fungi. Therefore, we repeated the experiment this time; both 8 hpi germlings and 14 hpi filaments were irradiated with 200 J/m^2^ UVC (Fig. 1B, *F. mangiferae*). Unlike at 14 hpi, we did not observe a significant induction of ribosome biogenesis at this developmental stage, although there is a slight shift towards disomes. A similar picture is also revealed for *F. oxysporum*; UV causes induction in ribosome biogenesis, although to a lesser extent than *F. mangiferae* (compare Fig. 1A with Fig. S3A).

The UV induction of ribosome biogenesis is observed at 14 hpi but not at the conidia stage (Fig. S3A and B).

### Translation capacity is induced in *Fusarium* species by UV irradiation at transcription and ribosome maturation levels

Ribosome biogenesis is the sum of several processes, the first of which is transcription of rRNA. To test whether rRNA is indeed induced at the transcriptional level by UV, we used Actinomycin D to inhibit RNA pol I activity (Perry and Kelley 1970, Schöfer, Weipoltshammer et al. 1996). 14 hpi filaments were exposed to 500 nM Actinomycin D two hours before UV irradiation with 200 J/m^2^. After irradiation, the fungi recovered in the presence of the inhibitor for 2 hours. Ribosomes were separated using a sucrose gradient as described above. Actinomycin D completely suppressed UV-induced ribosome biosynthesis. The effect of Actinomycin D was similar in both *F. mangiferae* and *F. oxysporum* (Fig. 2A, Fig. S4 correspondingly). Next, we asked whether the ability to form peptide bonds per se is increased following UV exposure. To this end, we harvested ribosomes from irradiated and non-irradiated cells at different time points after UV recovery and incubated them with biotinylated puromycin. Thus, translation of polypeptide chains can be detected using streptavidin-linked peroxidase (PUNCH-P; (Aviner, Geiger et al. 2013) and Materials and Methods). We observed a time-dependent increase in the puromycin-peptide signal following irradiation in *F. mangiferae* and to a lesser extent in *F. oxysporum* (Fig. 2B, Fig. S3C). We also tested the effect of Actinomycin D on UV-induced translation using the PUNCH-P protocol in *F. mangiferae*. We show that UV-induced translation is significantly delayed under Actinomycin D exposure but eventually occurs to a much lesser extent (Fig. 2B). This observation caused us to suspect that UV also induced ribosome maturation to some extent. To test this, we incubated irradiated cells with the ribosome maturation inhibitor Rbin2 for 14 hours. Filaments of *F. mangiferae* were treated with 1 µM Rbin2 for 1 hour before irradiation (200 J/m^2^) and then incubated with the inhibitor for an additional 2 hours. When we determined translation efficiency using the PUNCH-P approach under these conditions, the UV-induced translation capacity was suppressed with no recovery (Fig. 2B). The comparison between the dynamics of translation of the cells exposed to Actinomycin D and Rbin2 suggests that

**Figure 2.**
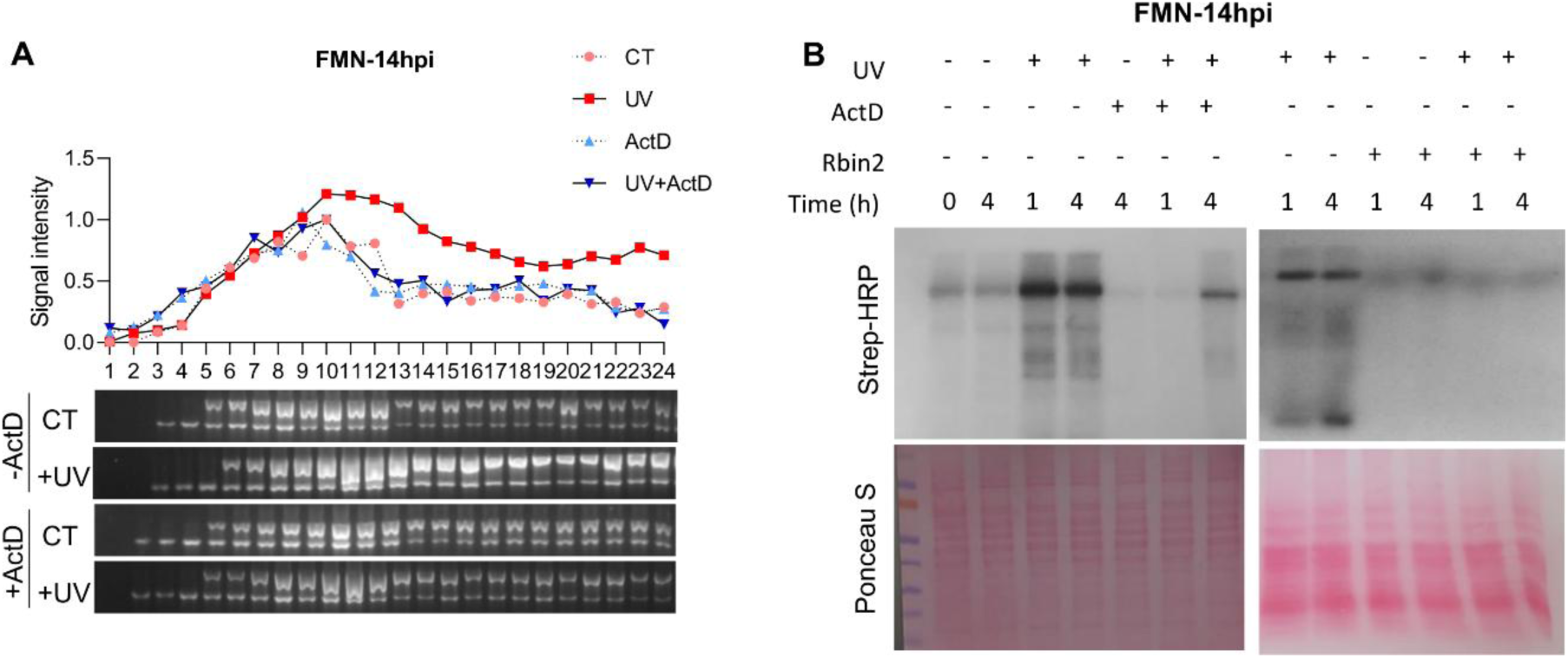
UV-induced translation capacity occurs at the ribosome transcription and maturation levels. (A) 14 hpi *F. mangiferae* (FMN) filaments were treated or not with Actinomycin D for 1 h and then irradiated with 200J/m^2^ UV. UV recovery occurred for 2 h in water at the same Actinomycin D concentrations. Polysome analysis was performed as described in the Materials and Methods section. (B) 14 h Fungal filaments were UV irradiated with 200 J/m^2^ or not, and then recovered for the indicated time points in water. Ribosomes were separated from the extracts using a sucrose cushion (see Methods). The purified ribosomes were incubated with Biotin-Puromycin to label nascent peptide chains (see Methods – PUNCH-P). Finally, the newly synthesized proteins were detected by western blotting with streptavidin-HRP and Ponceau S staining for total proteins. This procedure was followed in three different ways: 1; with no treatment at all, 2; with pretreatment of Actinomycin D as described in A, 3; with pretreatment of Rbin2 for 0.5 h, and then UV recovery in water at the same concentration for 2 h. The right panel of the western blot was cropped to fit into the figure; the original figure is shown in Fig. S5.

UV also induces ribosome maturation, and this is why, even under Actinomycin D exposure, minimal induced translation can be observed.

### Ribosome induction facilitates UV survival and repair of UV-induced DNA damage

Next, we wanted to examine whether inducing ribosome biogenesis and translation capacity improves UV survival. To this end, we exposed fungi to 500 nM Actinomycin D and then irradiated them. This time, the cultures were held in water for 24 hours after irradiation with or without Actinomycin D (liquid holding). In *F. mangiferae*, adding Actinomycin D increased the sensitivity of the fungi to 50 J/m^2^ of UV. The effect was observed in all the examined stages of fungal development, but was the strongest when 14 hpi filaments were irradiated (Fig 3A). Similar results were obtained when *F. mangiferae* cells were irradiated in the presence of 0.4 µg/ml cycloheximide, a protein synthesis inhibitor. Under these conditions, cycloheximide alone did not significantly reduce viability, but when combined with irradiation, viability decreased more than with irradiation alone. This phenomenon was observed in all stages, but the strongest decrease in viability (30%) occurred when 14 hpi filaments were irradiated (Fig 3B). Actinomycin D and Cycloheximide exposures were synthetic lethal with UV in F. oxysporum; however, there was no significant effect on the developmental stage (Fig. S6).

**Figure 3.**
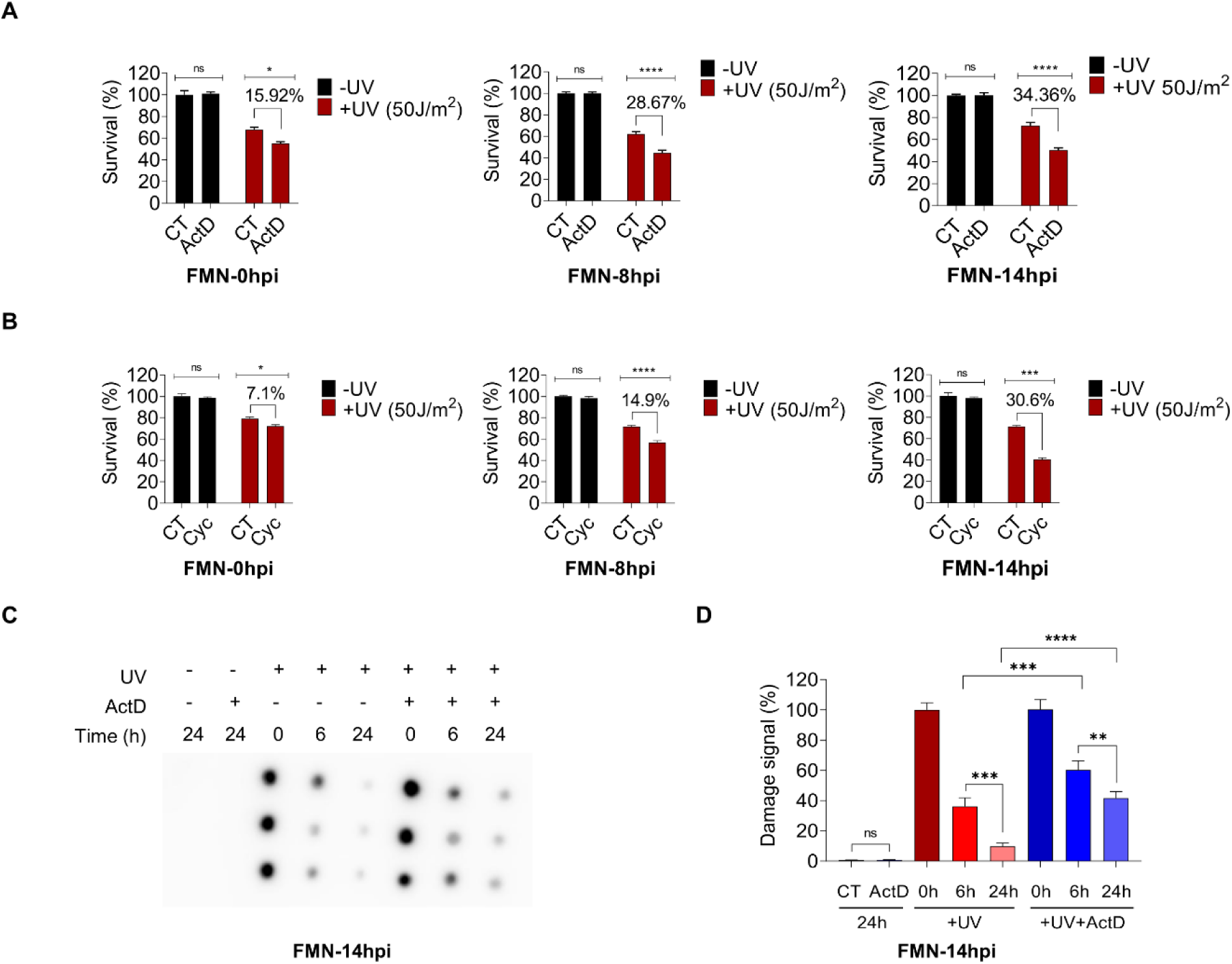
Blocking the UV-induced ribosome biogenesis and translation reduces UV survival and damage repair. *F. mangiferae (*FMN) conidia were inoculated for the indicated time points, then irradiated or not. (A) One hour before irradiation, the fungi were pretreated with or without Actinomycin D (ActD). After irradiation, the fungi were recovered for 12 h in water with or without Actinomycin D. (B) One hour before irradiation, the fungi were pretreated with Cycloheximide (Cyc) or not. After irradiation, the fungi were recovered in water for 6 h, with or without Cycloheximide. For both (A) and (B), after UV recovery, the fungi were plated on PDA. Survival was calculated as the ratio of the number of colonies that grew after exposure to Actinomycin D or Cycloheximide to the number of colonies that grew without such exposure (CT). 100% survival refers to the number of colonies without UV or chemical exposure. The results represent the average of three biological replicates. The experiment was repeated three times. Paired-wise, two-way ANOVA was used to calculate significance. *p<0.05, ****p<0.0001. (C-D) 14 hpi filaments were irradiated with 100 J/m^2^ and recovered in water with or without Actinomycin D for the indicated time points. After recovery, genomic DNA was isolated; UV lesions were detected by immuno-dot blot assay using antibodies against CPD (see Methods). (D) Quantification of damage signal from (C). The damage signal is shown relative to the initial damage inflicted. The damage signal at time point 0 after UV exposure was arbitrarily set to 1. The results are based on an average of six biological replicates, and error bars represent SD. Significance was calculated using Tukey’s multiple comparisons test, **p<0.05, ****p<0.0001.

Next, we wanted to know if UV repair is reduced in irradiated *F. mangiferae* filaments treated with Actinomycin D. In a liquid holding experiment of irradiated filaments for 24 hours, we were able to show that Actinomycin D reduced the repair of pyrimidine dimers. Without Actinomycin D, only 10% of the signal from the UV lesion remained, whereas with Actinomycin D, 40% remained (Fig. 3C&D). In summary, blocking UV-induced ribosome biogenesis sensitizes Fusarium to UV and reduces UV repair. However, the effect, while significant, is relatively modest and reflects a severe shutdown of the DNA damage response, as seen, for example, in the abolition of the SOS response.

### mRNAs related to gene expression and proteome health are associated with ribosomes following UV irradiation

Next, we sought to identify the genes whose translation is induced by UV. To this end, we performed an RNA-seq experiment of 14 hpi irradiated filaments two hours after irradiation. We sequenced both the total RNA population and higher- and lower-sucrose gradient fractions (fractions 3-12 and 13-24, respectively) (See Materials and Methods; all data are available in the GEO site under project GSE305365).

We first compared the number of differentially expressed genes by UV at two hours post-irradiation with that at earlier time points(Milo, Namawejje et al. 2024) and, as expected, the abundance decreased dramatically over time (Fig. S7). Also, the gene composition changes: the overlap between the induced genes within the first hour was much higher than at 2 hpi (Fig. S7). These findings suggest that, after two hours, the transcriptional response to UV is declining.

We identified mRNAs that are preferentially associated with ribosomes with and without UV irradiation. In support of our results, UV-irradiated fungi had more than triple the number of distinct mRNA species associated with polysomes (430 vs 130). Next, we identified the genes that are differentially associated with ribosomes only after UV irradiation and analyzed them by function. The genes can be divided into several groups. The largest group comprises genes important for gene expression: transcription, splicing, polyadenylation, ribosome biogenesis, and protein folding (Fig. 4). In this group, genes such as *ran1* were also found to facilitate the export of ribosomal subunits from the nucleus. This could be the reason why Rbin2 was more effective in suppressing UV-induced translation than Actinomycin D (Fig. 2). The second group of genes is related to metabolism, which can be further divided into mitochondria/electron chain transfer genes and amino acid metabolism (under the group of tryptophan biogenesis in Fig. 4, found genes that are related to the biosynthesis of other amino acids). The third group of genes is related to communication with the environment, either secretion or internalization. Among them, we identified genes related to the endoplasmic reticulum and coated pits. The fourth group seems most relevant to UV exposure, as it includes, among others, DNA repair and cell cycle genes. The automatic classification by the STRING algorithm of TC-NER (transcription-coupled repair) includes, in fact, genes of general NER. The most relevant gene in this group for UV repair is *DDB1*(Iovine, Iannella et al. 2011). Another enriched module is double-strand break repair by homologous recombination, which includes genes such as *rad55* and *mus81*, DNA synthesis (DNA polymerase delta, *psf1*, *mcm3*), and translesion DNA synthesis (DNA polymerase zeta and DNA polymerase kappa). Genes involved in cell cycle checkpoints, such as *cdc6* and the aurora kinase, were also identified (Fig. 4). The mRNA gene modules associated with lighter ribosomal fractions were similar to the heavy ones. They included genes related to translation, mitochondria, and the UV repair gene *DDB1* (Fig. S8). From all the above, it seems that two hours post-irradiation, the early induced transcripts are in the process of translation. Indeed, 165 of 430 genes associated with polysomes only after UV were induced by UV within the first hour (Fig. S9). This is not just because these genes are naturally associated with polysomes, because the proportion of these genes associated with un-irradiated polysomes is significantly lower (Fig. S9).

**Figure 4.**
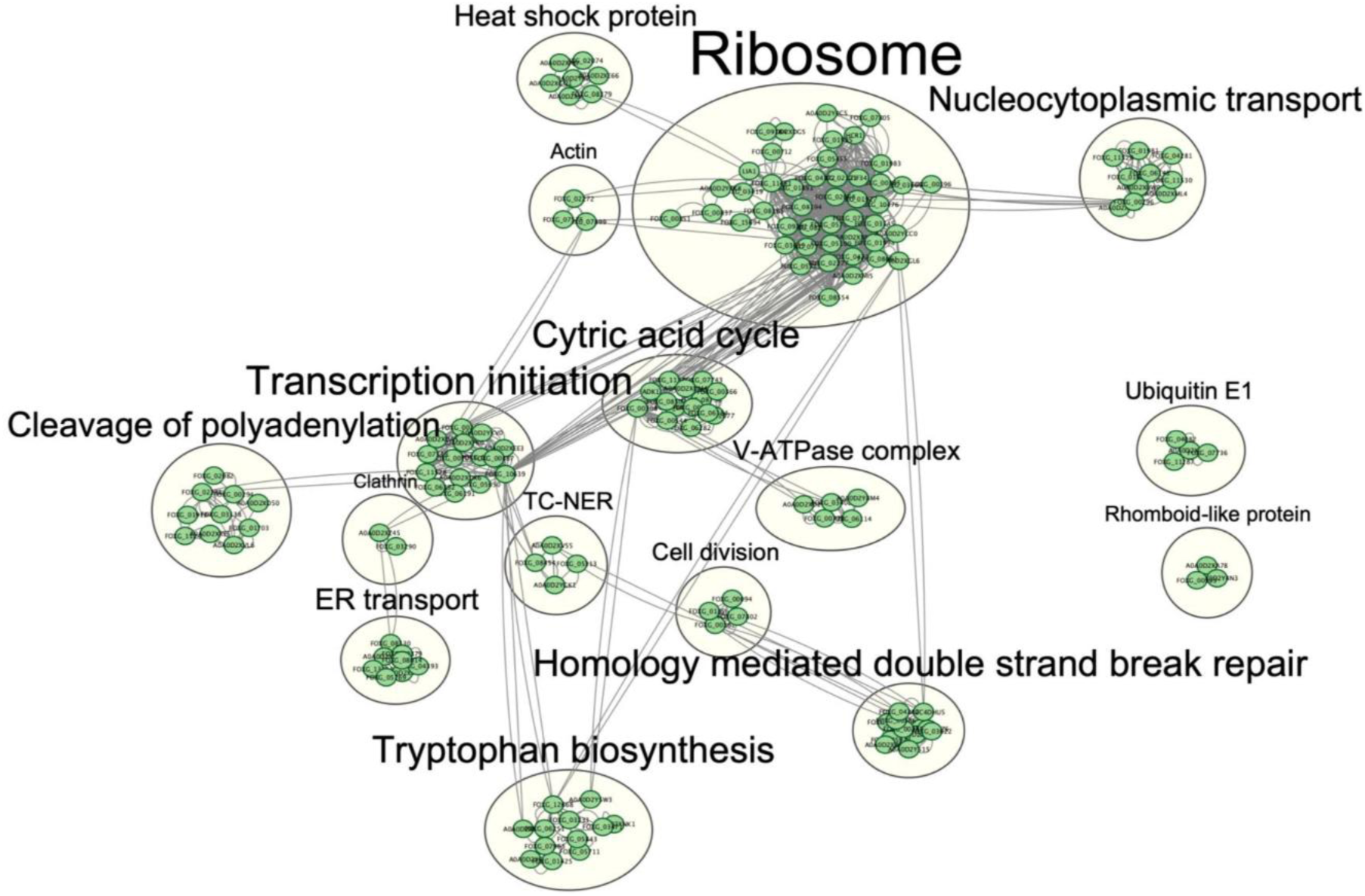
mRNAs associated with ribosomes after UV induction are clustered into gene expression and DNA damage response modules. 14 hpi filaments of *F. mangiferae* were irradiated or not with 200 J/m^2^. Two hours after recovery in water, ribosomes were separated by sucrose density gradient centrifugation as described in Fig. 1 and in the Methods section. RNA was purified from fractions 13-24, which represent polysomes, as described in the Methods. Next, the mRNA was further purified using poly dT and sent for sequencing. After sequencing, reads that mapped to ribosomal RNA units were removed from the analysis, as described in the Methods. To complete the data set, total mRNA was purified from the same cultures without further ribosome separation. Genes enriched in polysome fractions, with and without UV exposure, were identified first (raw data are available at GSE305365). Next, genes enriched only in irradiated polysomes were identified and analyzed using STRING to reveal UV-regulated translation modules.

### TOR is necessary for full UV-induction of ribosomes

Finally, we sought to understand the mechanism regulating UV-induced translation capacity. The TOR signaling pathway is known to regulate ribosome biogenesis by induction of transcription factors. Therefore, we tested whether inhibiting the TOR signaling pathway using rapamycin would suppress the UV induction of ribosome biogenesis and translation-related genes. To this end, we pretreated *F. mangiferae* filaments with 5 µg/ml rapamycin two hours before irradiation. We harvested the RNA 30 minutes post-irradiation and sequenced it. As a control, we harvested RNA from untreated filaments and from filaments treated with rapamycin but not irradiated.

We first tested how rapamycin affects UV-induced gene expression. We compared the genes induced by UV under rapamycin exposure (UV+rapamycin) to no treatment, as we determined (Milo, Namawejje et al. 2024). Pretreatment with rapamycin completely changes the effect of UV on gene expression. There was very little overlap between the genes that were induced by UV with or without rapamycin (Fig. S10A). Importantly, while UV induces ribosome biogenesis genes in the absence of rapamycin (Fig. S1(Milo, Namawejje et al. 2024)), we did not observe this effect when cells were exposed to rapamycin. However, UV induces gene expression even when filaments are pretreated with rapamycin. After removing the contribution of rapamycin to gene induction, about 1000 genes are induced by UV compared with untreated filaments (no UV, no rapamycin). Functional characterization of UV-induced genes during rapamycin exposure reveals carbon and amino acid metabolic modules (Fatty acid metabolism, pyruvate and ethanol metabolism) (Fig. S10B). Next, to directly test the effect of UV under rapamycin conditions, we compared gene expression under rapamycin exposure, with or without UV. Interestingly, here we do observe induction of ribosome biogenesis and translation-related genes (Fig. S11A). We also observed induction of genes related to nitrogen and amino acid metabolism. Several modules are downregulated by UV, including splicing and mitosis, but what we find most interesting are genes related to gluconeogenesis (Fig. S11B). To sum up, it seems that, with rapamycin pre-treatment, UV alters filament metabolism.

Next, we performed a UV survival experiment the same way we did with Actinomycin D and Cycloheximide. In *F. mangiferae*, rapamycin did not reduce UV survival at the filament stage (Fig. 5C, comparison of the effect of UV between 0 and 0.5 µg/ml rapamycin). Intrigued by these results, we measured rapamycin toxicity with and without UV irradiation. Surprisingly, UV exposure rescued rapamycin toxicity across all developmental stages, especially at the germling (8 hpi) stage (Fig. 5B). We tested rapamycin-UV interplay in *F. oxysporum* f. sp. *lycopersici* (*Fol*) 4287. This strain is resistant to rapamycin, likely due to a *tor1* gene duplication. We performed a 12-hour liquid-holding UV survival experiment with a range of rapamycin doses. We observed that UV improved rapamycin survival, at least in some of the developmental stages.

**Figure 5.**
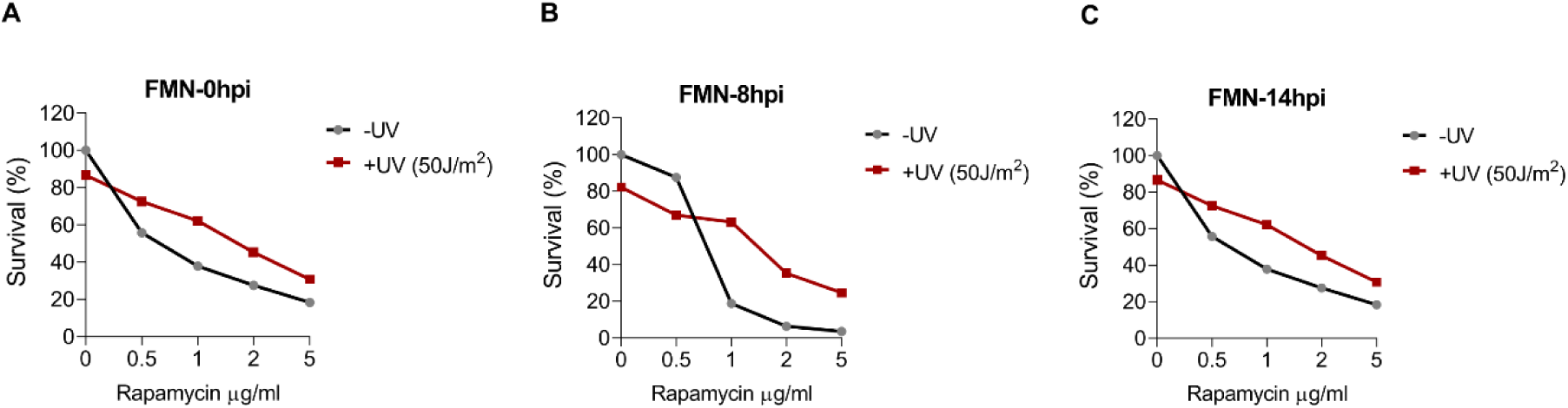
UV rescues *F. mangiferae* from rapamycin toxicity. (A). *F. mangiferae* were inoculated at the indicated time points in PDB, then incubated in water for 1 h with the indicated doses of Rapamycin. Next, the cultures were either irradiated or not and recovered in water containing the indicated doses of Rapamycin for 12 h. Cells were spread over PDA plates. Survival was calculated by dividing the number of colonies that grew after each treatment by the number of colonies in the culture that was not irradiated or treated with Rapamycin (considered as 100%)

Interestingly, the opposite was also true: rapamycin rescued UV toxicity in all tested stages of fungal development (Fig. S12). This means that there is apparent positive feedback between rapamycin and UV.

## Discussion

Here, we show that UV induces translation in two Fusarium species (Fig. 1, 2, Fig. S3,4). This contrasts with many reports on the negative effects of DNA damage and other stresses on ribosome biogenesis. The question is why? Unlike the SOS response, our results do not support the induction of many DNA repair genes (Maslowska, Makiela-Dzbenska et al. 2019) (Fig. 4, Fig. S8). We observed a positive and significant effect of ribosome biogenesis on UV survival and DNA repair, but it is not comparable to the effect of the SOS pathway in bacteria (Fig. 3 and (Keller, Overbeck-Carrick et al. 2001))). Several DNA repair genes are specifically associated with ribosomes following UV (Fig. 4, Fig. S8). Still, there are significantly higher mRNAs that belong to transcription, translation, protein secretion, and other functions that are associated with ribosomes following UV (Fig. 4, Fig. S8). We think that the UV induction in translation capacity is to compensate for the negative effect of UV on transcription and, consequently, translation (Nieto Moreno, Olthof et al. 2023). As a support mechanism, the association of mRNA encoding housekeeping proteins, including chaperones, with ribosomes is enriched following UV exposure to maintain homeostasis. We are currently testing whether other ways to block mRNA transcription elicit the same response, or whether the DNA damage response strictly mediates it.

Why was UV induction of ribosomes never observed in other organisms? We suggest it has to do with the (in)ability of *Fusarium* species to overcome UV effects on transcription. In many organisms, from *E. coli* to humans, transcription-coupled repair is dedicated to overcoming DNA damage-induced stalling of transcription (Gaul and Svejstrup 2021). We hypothesize that transcription-coupled repair in *Fusarium* species is less efficient than in other organisms and, consequently, that transcription resumption following DNA damage is slower. In support, we have previously reported that mRNA levels of components of the basic mRNA transcription machinery are induced by DNA damage (Milo-Cochavi, Pareek et al. 2019). Here we observed that the association of these mRNAs with ribosomes is increased following UV exposure (Fig. 4). This could be the response in *Fusarium* for the destruction of stalled transcription apparatus in case transcription-coupled repair is inefficient, or it could be that normal transcription machinery is not enough to trigger TCR and more transcription initiation is needed. It was previously demonstrated that waves of RNA polymerase II are released from pause to transcription elongation following UV exposure, in what appears to be a mechanism to reduce mutagenesis (Lavigne, Konstantopoulos et al. 2017).

Another explanation for why we observed UV-induced ribosome biogenesis in Fusarium but not in other species may be the developmental biology of filamentous fungi. UV-induced ribosome biogenesis is observed at 14 hpi when the filaments are formed (far less in conidia or germlings) (Fig. 1, Fig S3). While growth at 14 hpi is still fast, most of the nuclei in the filament are already dormant. Although most nuclei should be mitotically dormant, among the genes associated with polysomes following UV are those related to the cell cycle, specifically mitosis. This may suggest that different nuclei translate distinct sets of genes following UV exposure, a heterogeneity absent in *E. coli* or yeast cells. One aspect to consider is the division of labor and the interplay between TOR signaling and PKA in ribosome biogenesis ) Guerra, Vuillemenot et al. 2022, Plank 2022(. The balance between these two pathways may change during fungal development in ways not observed in organisms studied to date. *Fusarium mangiferae* is more sensitive to rapamycin at the germling stage than the filament stage (Fig. 5). It is possible that when TOR signaling is dominant, UV induction is insignificant, but when the balance is changed towards PKA, either by TOR inhibition or during development, UV induction is more significant. It is possible that UV-induced PKA signaling activates pathways that, in turn, activate TOR, which is ultimately required for UV-induced ribosome biogenesis (Fig. S10).

The results presented here pose many questions. We still do not know which molecular switch is activated during filament development and facilitates UV-induced translation. There are differences in the UV response between *F. mangiferae* and *F. oxysporum*; some of these may be due to differences in the timing of activation of this switch post-inoculation. We hypothesize that both the developmental switch and the differences between the fungi stem from modulation of crosstalk between PKA and TOR signaling.

Translation requires a lot of energy. In agreement, mitochondrial genes are induced at the transcriptional and ribosome-association levels following UV exposure (Fig. S1, Fig. 4, Fig. S8 (Milo-Cochavi, Adar et al. 2019, Milo, Namawejje et al. 2024)). What are the metabolic adjustments that the fungi need to make to enable this energy production?

When TOR is repressed by rapamycin, many of the differentially expressed genes are involved in metabolism (Fig. S10B, 11, Tables S2 and S3 and S4). Could it be that UV induces metabolic rewiring that increases the rate of energy production? And if so, how? Finally, from a plant pathology standpoint, the commitment to ribosome biogenesis is so high that one can speculate that filaments following UV exposure are more vulnerable to stress. Can this be used to fight *Fusarium* pathogens?

## Materials and Methods

### Fungal isolates and culture conditions

Spores of *F. mangiferae* (strain MRC7560, isolated in Israel) and *F. oxysporum* (*F. oxysporum* f.sp. *lycopersici* strain 4278) were incubated in KNO3-based medium (KNO3: 10.1g, Sucrose: 30g, YNB (Yeast Nitrogen Base): 1.7g in 1L distilled water) on an orbital shaker (250 rpm) at 28 °C for 7 days. Microconidia were obtained by filtering the culture through a nylon cell strainer (40 µm mesh, Corning, USA) and resuspended in sterile water. Conidia were then inoculated into potato dextrose broth (PDB; BD Sparks, USA) at 1x10^8^ conidia/ml. The cultures were incubated on an orbital shaker at 28 °C for 8 or 14 h, then harvested immediately or treated as specified.

### UV irradiation

Spores or germlings of *F. mangiferae* or *F. oxysporum* (0, 8, and 14 h post-inoculation) were obtained by filtering 100 ml of the cultures. Suspensions containing 2 x 10^7 conidia/ml were irradiated with different UVC doses ranging from 30 to 200 J/m^2^ (254 nm, 15 W lamp, Osram, Germany); the incident fluence was measured with a radiometer (YK-35UV, Digital Instruments, USA). Irradiation dose and incubation time after irradiation were determined in downstream experiments (RNA sequencing, liquid holding, immunodot blot, polysome profiling, or PUNCH-P). Samples were harvested, immediately flash-frozen in liquid N2, and kept at −80 °C until further experiments.

### Ribosome isolation

Frozen samples were ground into a fine powder using a mortar and pestle in liquid nitrogen. Cell lysates were prepared in a cold lysis buffer containing 50 mM Tris-HCl (pH 7.5), 50 mM KCl, 10 mM MgCl_2_, 1% protease inhibitor cocktail (Roche) supplied with cycloheximide (10µg/ml, final concentration) homogenized with glass beads using Mini bead beater following by two cold centrifugations, the first one at 5000rpm, 4°C for 5 minutes and the second one at 12000 rpm for 30 minutes. The lysate supernatant was then transferred into ultra-centrifuge tubes, and 5 mL of 32% sucrose (in lysis buffer) was carefully added using the long-range needle. Equal amounts of nucleic acids (25 A260 units) were loaded. Samples were spun in a Beckman Coulter 70.1 Ti rotor at 50.000 rpm for 16 hours at 4°C. After discarding the supernatant, the resulting ribosomes, forming a clear pellet, were resuspended in 200μl extraction buffer (ribosome recovery) (Mehta, Woo et al. 2012).

### Polysome fractionation and RNA extraction

Polysome fractionation was carried out following the ribosomal isolation (dissolved pellet, above). 200 μl of the dissolved pellet was loaded onto a 12 ml linear sucrose gradient (10-50%) prepared in lysis buffer. The sucrose gradient was prepared using a gradient maker (Gradient Master™ from BioComp) according to the manufacturer’s instructions. The loaded samples were then centrifuged at 39,000rpm for 3 h at 4°C using an SW 41Ti rotor. Individual fractions were collected with a fraction collector. RNA was precipitated from 0.5 ml fractions by mixing each fraction with an equal volume of 6 M guanidine thiocyanate and 2 volumes of 100% ethanol, then incubating overnight at −20°C. The RNA was pelleted, washed, and briefly resuspended: it was pelleted in 75% ethanol by centrifugation for 30 minutes at 10,000g at 4 °C, and the supernatant was discarded. Then the pellet was washed using the same procedure with some differences. 0.5–1 ml of 75% ethanol was added to the pellet, which was then vortexed for a few seconds. Next, the pellets were incubated for 10–15 minutes at room temperature to dissolve any residual traces of guanidinium, then centrifuged for 5 minutes at 10,000g at 4 °C, with the supernatant discarded. Then the pellet was air-dried. Finally, the pellet was dissolved in 100 µl of either DEPC-treated water or freshly deionized formamide (For Formaldehyde gel electrophoresis). By incubating for 10–15 minutes at 60 °C to ensure complete solubilization. The samples were run by formaldehyde agarose gel electrophoresis (1.5% agarose, 5% formaldehyde in 1 × MOPS buffer (20 mM 3-[N-morpholino] propane-sulfonic acid (MOPS, M3183, Sigma), 5 mM sodium acetate, 1 mM EDTA)). 4.7 μl of RNA was mixed with 11.3 μl sample buffer (for 1.13ml: formamide 660 μl, MOPS buffer 10X 200 μl, formaldehyde 37% 270 μl), then heated to 60°C for 5 minutes, cooled on ice, and 4 μl loading dye was added. Gels are run for 1.5-2h in an ice bucket. The gels were stained with SYBR Gold (Invitrogen, S11494) by the standard procedure. ImageJ software was used to determine band intensities and, consequently, the ribosome signal. Then, for each gel image, the relative intensities of the different gradient fractions were determined. Next, the fractions with the highest intensities from each gel were run together on a single gel. The signal intensity was measured using ImageJ and assigned a relative intensity value compared with the unirradiated and untreated sample (signal intensity as 1). Finally, the relative intensities of all fractions across all gels were recalculated using the control sample value.

For RNA-seq experiments, fractions from 3-12 and 13-24 were combined and processed for RNA extraction using a Trizol LS 3:1 ratio according to the manufacturer’s instructions. After RNA extraction, the workflow followed the general next-generation sequencing protocol.

### Transcriptome analysis

Raw reads were trimmed using Fastp 38 and assessed for quality using FastQC (https://www.bioinformatics.babraham.ac.uk/projects/fastqc/). Ribosomal RNA gene sequences of *Fusarium fujikuroi* (EF1) were downloaded from ENSEMBL Fungi Biomart (http://fungi.ensembl.org/index.html) and used to construct a reference database for FMN. Raw reads were mapped against the reference rRNA database using HISAT2 39. Mapped reads were discarded, whereas unmapped reads were collected and remapped against the *F. mangiferae* [FMN] MRC7560 [GCA_900044065.1] reference genome. Mapped reads were then counted with HTSeq-count (Chen 2023). The reads were assembled and normalized, and gene expression levels were quantified using the DESeq2 R package (Zhang, Park et al. 2021). Differential expression was assessed for genes expressed in all combinations of UVC treatments vs. untreated controls. DESeq2 was used to identify differentially expressed genes using the negative binomial distribution, with adjusted P-values (P < 0.05) and log2 fold changes> +1 or < −1. To identify significantly enriched genes in the ribosomal fraction only after UV exposure, ribosomal-enriched genes with and without UV were analyzed using a Venn diagram in Venny (https://bioinfogp.cnb.csic.es/tools/venny/).

### Functional enrichment and network analysis

Functional enrichment analysis was performed using the gprofiler2 v0.2.3 R package 40 and one-tailed Fisher’s exact test. The g: SCS method was used to compute multiple-testing correction, with P < 0.05 as the significance threshold. (50). To reveal the genes that are uniquely expressed under distinct conditions, either up- or downregulated differentially expressed genes from different polysome fractions were intersected based on condition (treated vs. untreated). Differences in each set (unique in treated or unique in untreated) were analyzed using the protein-protein interaction network analysis in the STRING database (Kolberg, Raudvere et al. 2023) and visualized in Cytoscape v3.9.1 (https://cytoscape.org/index.html). Clusters were identified and annotated using the Cytoscape AutoAnnotate plugin v1.3.5 (http://baderlab.org/Software/AutoAnnotate).

### PUNCH-P and Western Blot

The PUNCH-P protocol was performed as described in (Aviner, Geiger et al. 2013). 5 A260 units of Ribosomal proteins were used for each labeling with Biotin-Puromycin. 100 pmol of Biotin-dC Puromycin (Jena Biosciences cat# NU-925-BIO-L) was added per 1 OD260 unit of ribosome-containing solution. As a control, non-puromycylated (-P) reactions were also used. The reactions were incubated at 37°C for 30 minutes, after which the proteins were precipitated with chloroform and methanol to remove excess Puromycin for subsequent SDS-PAGE. Briefly, after the reaction, water and 6x SDS buffer containing bromophenol blue were added to a final volume of 150 µl. Following the general protocol for protein precipitation with 4 volumes of methanol, 2 volumes of chloroform, and 3 volumes of water. The pellet will have some blue color from the Bromophenol blue for easy recognition. The protein pellet was then resuspended in SDS buffer and loaded onto a 12% SDS PAGE. Proteins were transferred onto an Amersham Protran 0.45 μm nitrocellulose membrane (GE Healthcare, 10600003) at 25V overnight using the wet transfer method. The membrane was stained with Ponceau S solution (2% solution in 2% acetic acid) to visualize transfer efficiency. The membrane was washed off this stain with water for 5 minutes, followed by a 5-minute wash with TBST (Tris-buffered saline with 1% Tween-20). The membrane was blocked with 5% skim milk in TBST for 1 hour at room temperature. After blocking, the membrane was washed three times, each for 5 minutes at room temperature in TBST, to remove any remaining skim milk solution. The membrane was then incubated with a 1:5000 dilution of streptavidin-horseradish peroxidase (HRP) (Sigma SA10001) in TBST for one hour at room temperature.

### Liquid holding assays

Conidia, germling or filaments were kept in distilled water added with Actinomycin D (Sigma, A9415) (500 nM or as described in the figures), Rapamycin (Thermofisher Scientific, J62473.MF), Rbin2 (Sigma, 2032282) (1 µM), Cycloheximide (Sigma, 01810) (0.3 µg/ml) and held for different incubation times before plating on PDA plates.

Colonies were counted after 2 to 3-d post-inoculation and the suitable concentration with the best holding time which showed 100% survival were chosen for further experiment. After UV-irradiation, conidia or germling were kept in distilled water added with desirable doses of Actinomycin D, Rapamycin, Rbin2, Cycloheximide and held for the indicated time to check the effect of drugs on the survival under UV. Colonies were counted after 2 to 3-d inoculation and the additive effect would be evaluated. At least three sets of experiment were repeated, each with three biological replications. The same procedure was used also to test the effect of the different inhibitors on ribosome biogenesis and translation capacity (PUNCH-P protocol).

### Immunodot blot assay

After UV irradiation, samples were kept in distilled water containing Actinomycin D for the indicated time (6 to 24 h), then harvested and processed for DNA extraction using the CTAB-based method. In short, samples were ground into a fine powder using liquid nitrogen and disrupted with a Minilys bead beater for 60 s at medium speed in RLC buffer. Genomic DNA was purified using 2% (wt/vol) hexadecyltrimethylammonium bromide (CTAB) buffer, followed by a chloroform-isoamyl alcohol (24:1 [vol/vol]) phase separation procedure and precipitation in a final concentration of 50% ice-cold isopropanol. An immunodot blot assay was used to quantify DNA lesions, as described previously. Briefly, after denaturation at 95°C for 10 minutes, 400 ng of DNA was combined with an equal volume of 2 M ammonium acetate and placed on ice. Each DNA sample was spotted onto a nitrocellulose membrane (soaked in 1 M ammonium acetate buffer for 10 minutes at room temperature), using a Bio-Rad dot blot manifold. The membrane was washed twice with 1 M ammonium acetate buffer and once in 6X saline sodium citrate buffer. Membranes were dried in a vacuum gel dryer (model 583; Bio-Rad, USA) for 90 minutes at 80°C. After blocking in 5% powdered milk in PBST (1 ml Tween 20, 100 ml of 10× phosphate-buffered saline [Biological Industries, Israel], 899 ml double-distilled water), membranes were probed with mouse anti-CPD antibody (NMDND001; Cosmo-Bio, Japan). Following secondary antibody application (peroxidase-conjugated; Jackson Immuno Research Laboratories, USA), enhanced chemiluminescence was used to detect the antibody-dependent signal from each DNA spot on film. The intensity of each spot was quantified using ImageJ.

## Acknowledgments

We thank Dr. Moshe Peretz for his expertise, guidance and support in ribosome profiling experiments. This work was supported by Israel Science Foundation Grant # 410/21

