## Supplemental Figures for "UV-induced translation capacity is regulated developmentally in fungi"

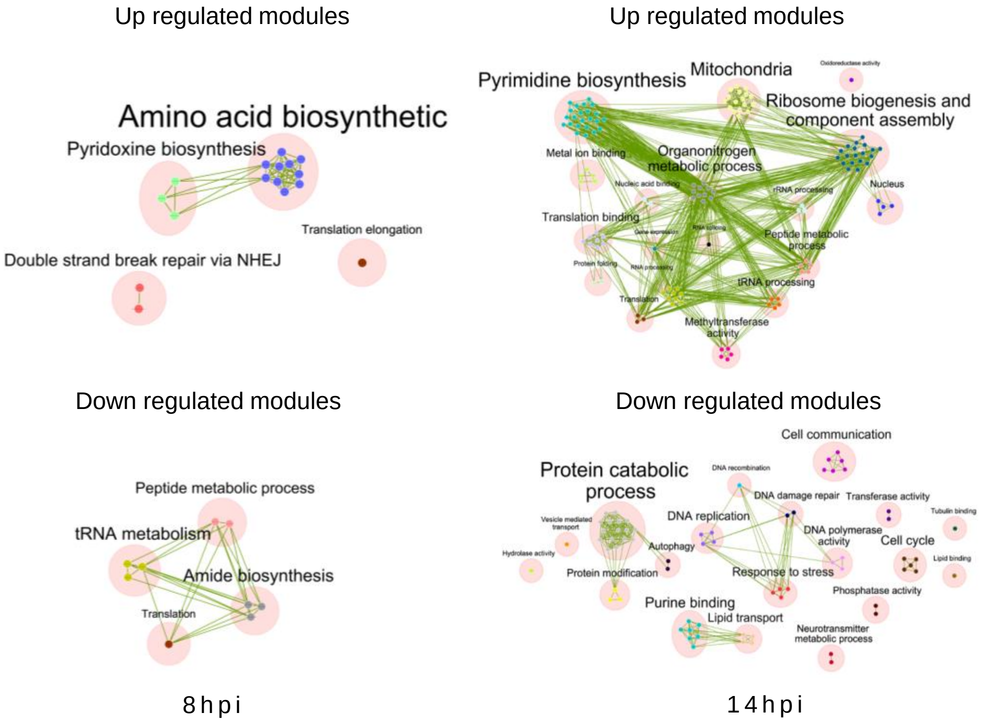

Up regulated modules
Amino acid biosynthetic
Pyridoxine biosynthesis
Translation elongation
Double strand break repair via NHEJ
Up regulated modules
Pyrimidine biosynthesis
Mitochondria
Ribosome biogenesis and component assembly
Metal ion binding
Organonitrogen metabolic process
Nucleus
Translation binding
Peptide metabolic process
tRNA processing
Translation
Methyltransferase activity
Down regulated modules
Peptide metabolic process
tRNA metabolism
Amide biosynthesis
Translation
Down regulated modules
Cell communication
Protein catabolic process
DNA damage repair
Transferase activity
DNA replication
DNA polymerase activity
Cell cycle
Autophagy
Protein modification
Response to stress
Phosphatase activity
Purine binding
Lipid transport
Neurotransmitter metabolic process
8 h p i
1 4 h p i

**Figure S1. Modules related to translation are induced by UV in *F. mangiferae* when filaments but not germlings are irradiated.**

Conidia of *F. mangiferae* were inoculated for 8 or 14 h in PDB medium. The germlings or filaments were harvested, re-suspended in water, and irradiated with 200 J/m2 after recovery for 0, 30, or 60 minutes in complete darkness. The mRNA was purified and sequenced; the raw data are found in GSE254119. The UV-differentially expressed genes across all time points were pooled and analyzed using STRING to identify up- and down-regulated modules.

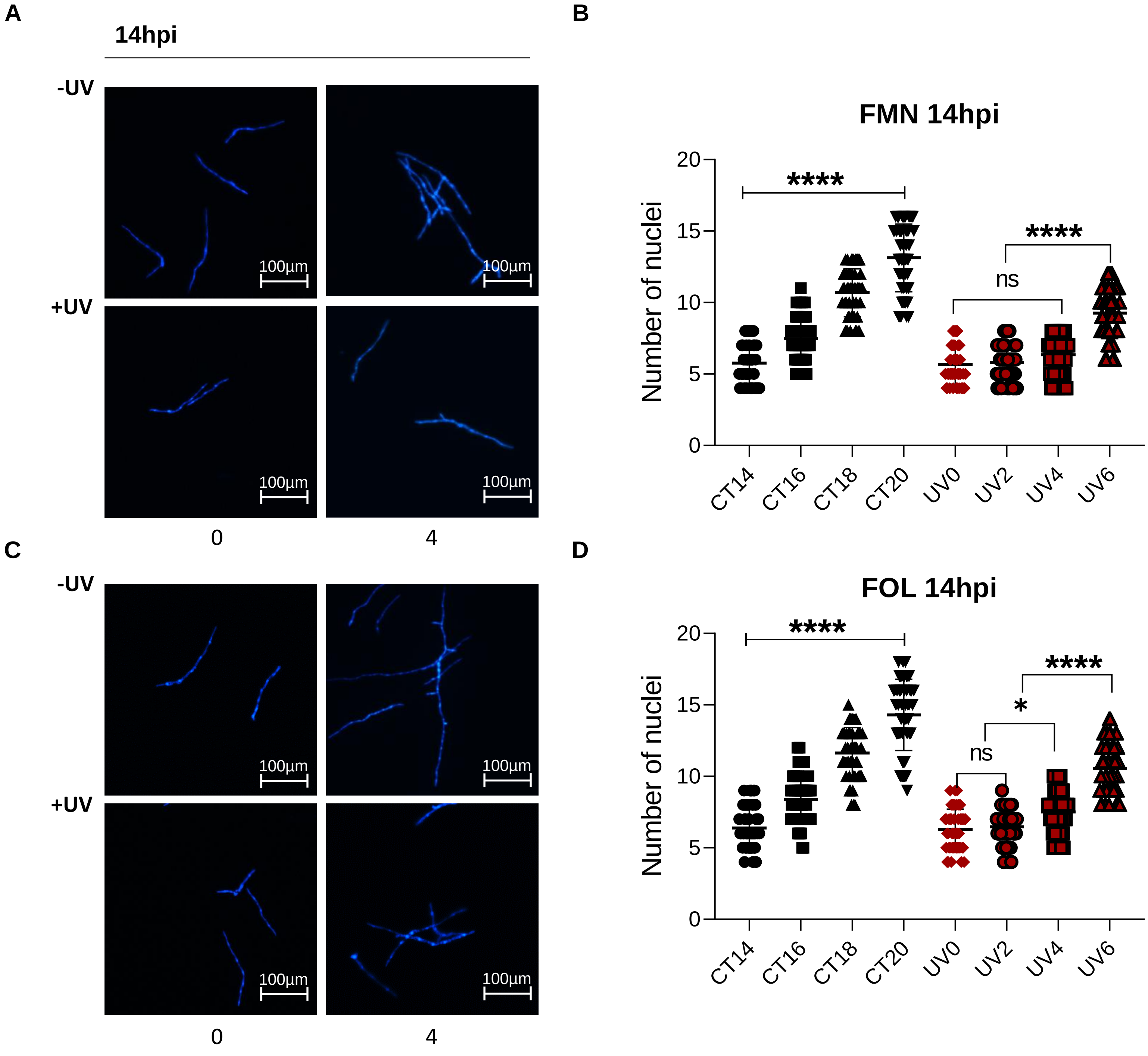

A
14hpi
-UV
+UV
100µm
0
4
B
FMN 14hpi
Number of nuclei
0
5
10
15
20
****
****
ns
CT14
CT16
CT18
CT20
UV0
UV2
UV4
UV6
C
-UV
+UV
100µm
0
4
D
FOL 14hpi
Number of nuclei
0
5
10
15
20
****
****
*
ns
CT14
CT16
CT18
CT20
UV0
UV2
UV4
UV6

**Figure S2. UV causes nuclear division delay in filaments of *F. mangiferae* and *F. oxysporum***

Representative images of *F. mangiferae* (FMN) (A) and *F. oxysporum* (FOL) (C), 14 hpi filaments were irradiated with 200 $J/m^2$ UV and then were released for two to six hours in PDB. Fluorescence indicating nuclear fragmentation by DAPI staining. After UV exposure, fungal cells were fixed with 4% paraformaldehyde in phosphate-buffered saline (PBS) for one hour at room temperature. Then, the filaments were washed before dehydrated in 75% ethanol overnight. Samples then were washed twice with PBS before DAPI staining (1 μg/ml). Scale bar, 100µm. (B-D) Distribution of nucleus numbers in filaments of *F. mangiferae* (B) and *F. oxysporum* (D). Significance was calculated using one-way ANOVA, Tukey test, * $p<0.05$, **** $p<0.0001$

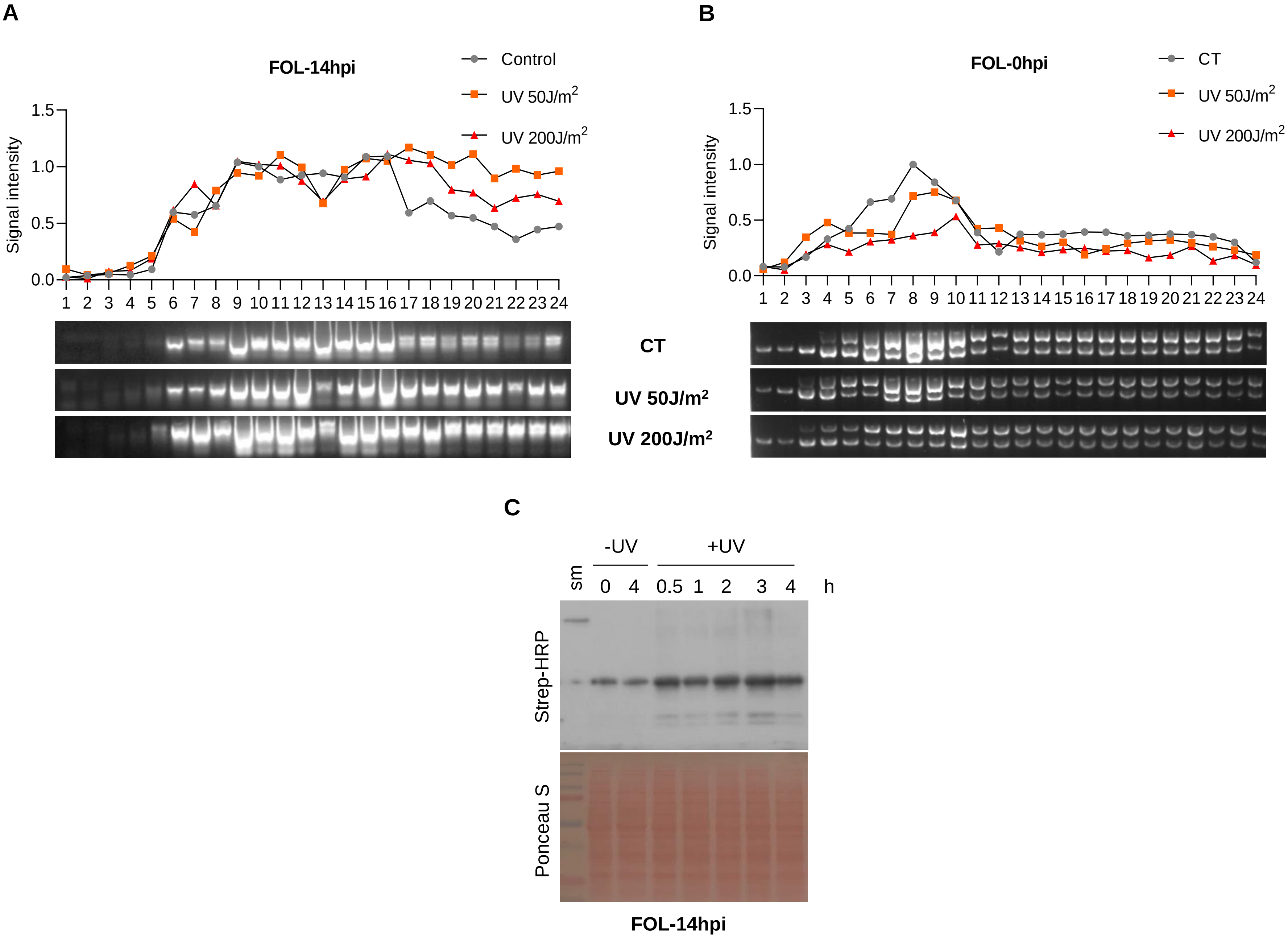
A
FOL-14hpi
Control
UV 50J/m2
UV 200J/m2
Signal intensity
1.5
1.0
0.5
0.0
1 2 3 4 5 6 7 8 9 10 11 12 13 14 15 16 17 18 19 20 21 22 23 24
CT
UV 50J/m2
UV 200J/m2
B
FOL-0hpi
CT
UV 50J/m2
UV 200J/m2
Signal intensity
1.5
1.0
0.5
0.0
1 2 3 4 5 6 7 8 9 10 11 12 13 14 15 16 17 18 19 20 21 22 23 24
C
-UV
+UV
sm
0 4 0.5 1 2 3 4 h
Strep-HRP
Ponceau S
FOL-14hpi

**Figure S3. UV induces ribosome biogenesis and translation capacity in filaments of *F. oxysporum***

Polysome gradient profiles of *F. oxysporum* (FOL)14 hpi (A) and microconidia 0 hpi (B) irradiated with 50 and 200J/$m^2$ UV doses. The RNAs from each fractions of polysome gradient were separated on 2% agarose gels; ribosome intensity was calculated based on the strongest band signal in the control sample (signal intensity as 1) (similarly to what is described in the legend of Fig. 1). (C) The effect of UV on translation capacity. Ribosome of *F. oxysporum* were isolated after UV exposure of 14 hpi filaments. Next, the ribosomes were labeled by Biotin-Puromycin for the indicated times and under the indicated conditions and detected by western blotting using streptavidin-HRP (similarly to what is described in the legend of Fig. 2B).

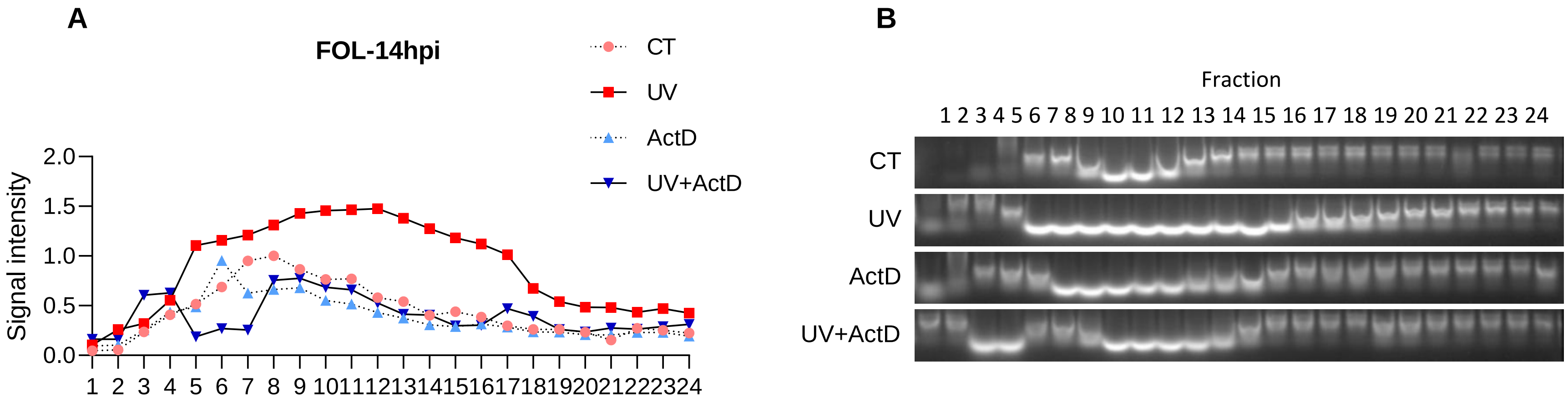


**Figure S4. Actinomycin D inhibits the UV-induced ribosome biogenesis in *F. oxysporum***

(A,B). Polysome gradient profiles of *F. oxysporum* (FOL) 14 hpi treated with 500 nM of Actinomycin and 200J/m$^2$ UV as described in the legend of Fig. 2A. The RNAs from each fractions of polysome gradient were separated on 2% agarose gels and the ribosome intensity was calculated based on the strongest band signal in the control sample (signal intensity as 1).

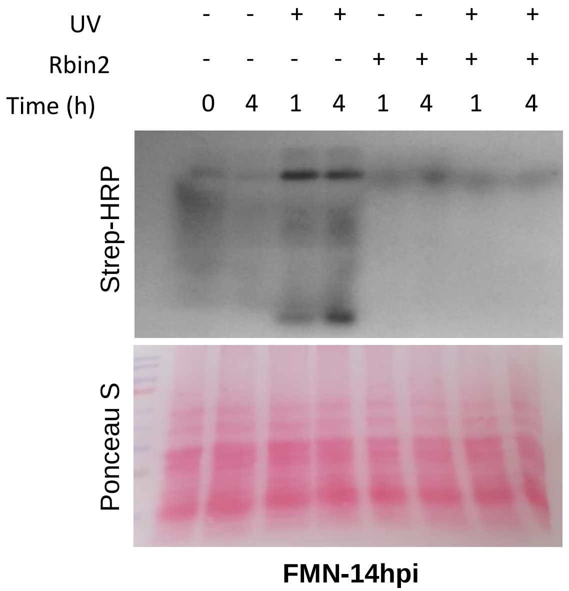


**Figure S5. Rbin2 suppresses UV-induced translational capacity in *F. mangiferae* (FMN)**

This is the full image of the cropped image presented in Fig. 2

**A**

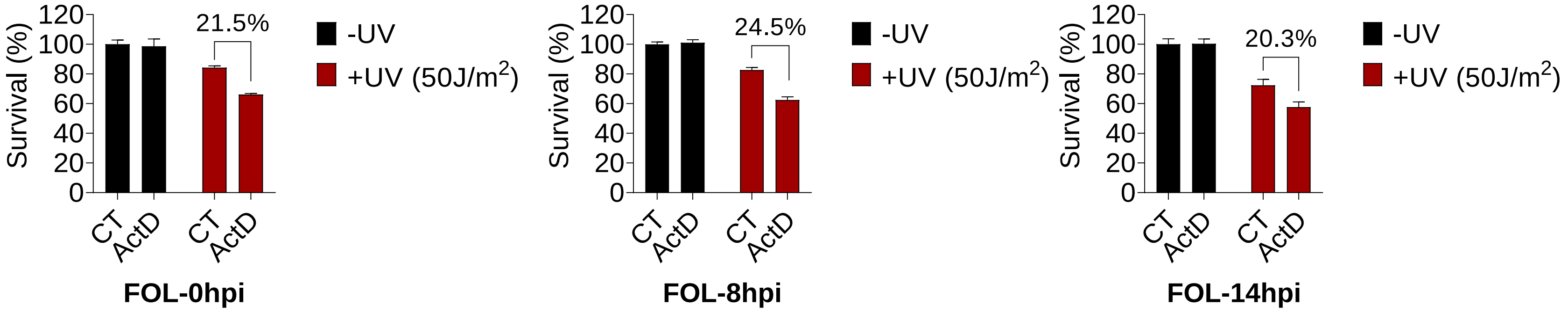
Survival (%)
120
100
80
60
40
20
0
21.5%
CT
ActD
CT
ActD
FOL-0hpi
-UV
+UV (50J/m2)
24.5%
FOL-8hpi
20.3%
FOL-14hpi

**B**

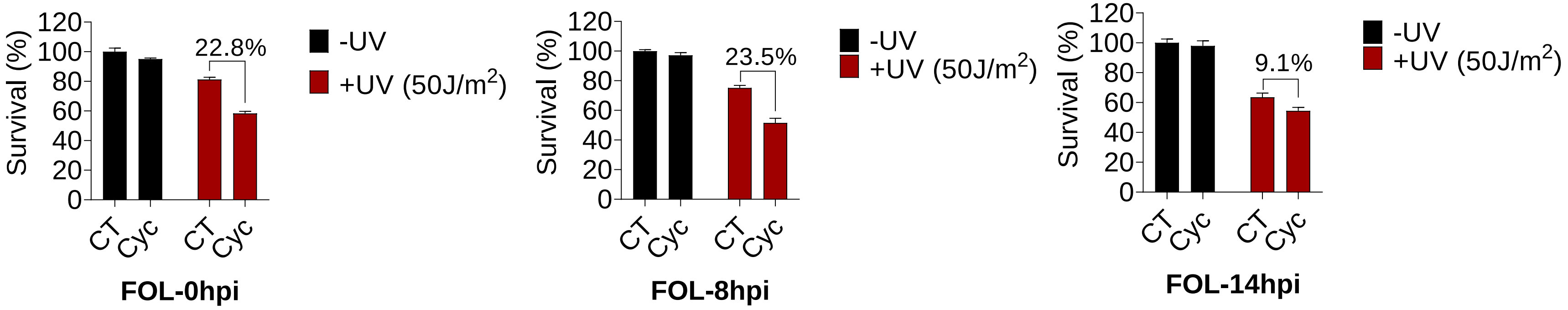
Survival (%)
120
100
80
60
40
20
0
22.8%
CT
Cyc
CT
Cyc
FOL-0hpi
-UV
+UV (50J/m2)
23.5%
FOL-8hpi
9.1%
FOL-14hpi

**Figure S6. Actinomycin D and Cycloheximide are synthetic lethal with UV in *F. oxysporum***

Liquid holding assay with 50J/m$^2$ UV dose and 500 nM Actinomycin D (A) and 0.3µg/ml Cycloheximide (B) of *F. oxysporum* (FOL) at different stage of development. Survival was calculated by dividing the number of colonies that grew after UV irradiation with the one without. The experiment was conducted similarly to what is described in the legend of Fig. 3A&B. The results represent the average and SD of 3 biological repeats. Significance was calculated using Paired wise, Two-way ANOVA, Tukey test. * $p<0.05$, ** $p<0.001$, *** $p<0.0001$

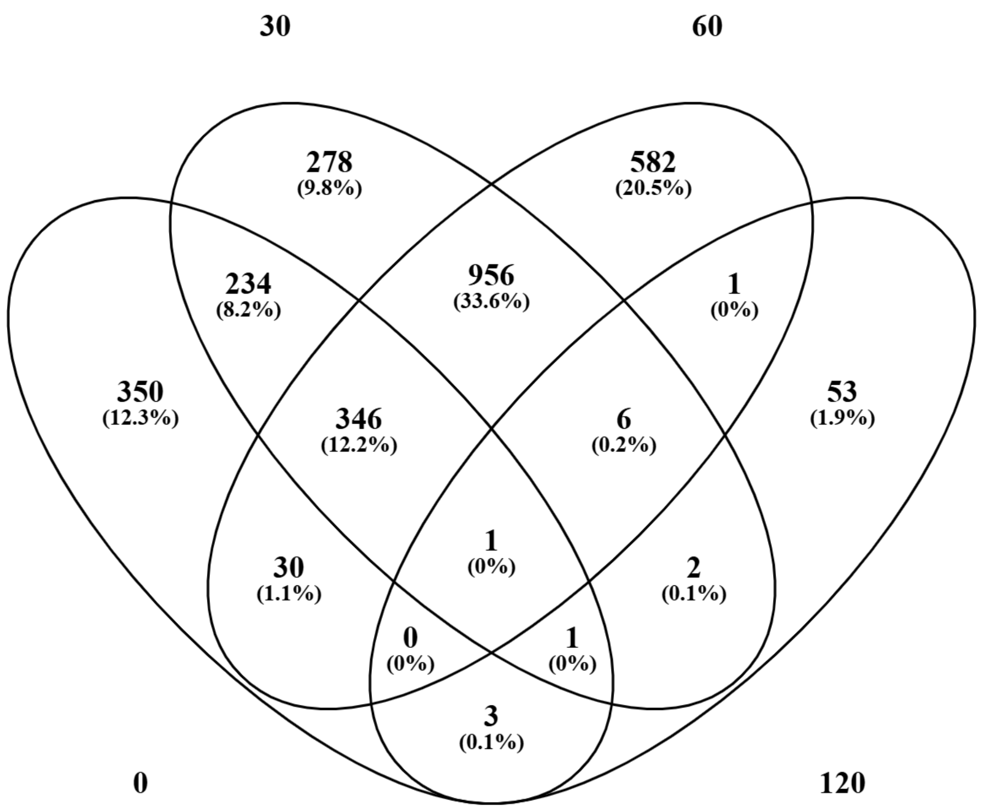

30
60
278
(9.8%)
582
(20.5%)
234
(8.2%)
956
(33.6%)
1
(0%)
350
(12.3%)
346
(12.2%)
6
(0.2%)
53
(1.9%)
1
(0%)
30
(1.1%)
2
(0.1%)
0
(0%)
1
(0%)
3
(0.1%)
0
120

**Figure S7. Delayed response of *F. mangiferae* filaments to UV: the early response differs in the number of induced genes and in gene composition.**

The number of genes induced by UV after irradiation of filaments. The numbers for 0, 30, 60 and 120 minutes after irradiation recovery times are shown. The data for the early time points (0-60) are taken from Milo et al.; Fungal Biology 2024 (see also the legend to Fig. S1). The data for the later time point (120 min) were collected as described in Materials and Methods and in the legend to Fig. 4. The overlap between the gene lists was created by Venny 2.1.

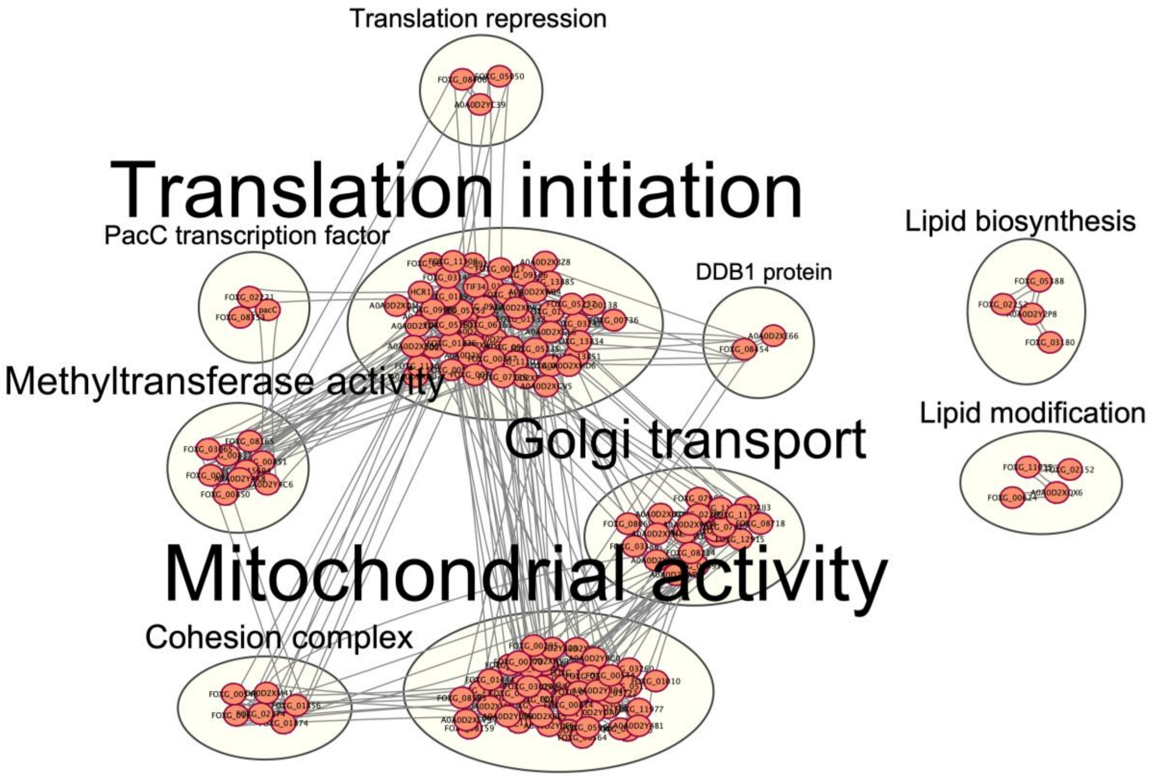
Translation repression
Translation initiation
PacC transcription factor
Lipid biosynthesis
DDB1 protein
Methyltransferase activity
Golgi transport
Lipid modification
Mitochondrial activity
Cohesion complex

**Figure S8. mRNA associated with ribosomes after UV induction are clustered to gene expression and DNA damage response modules.**

14 hpi filaments of *F. mangiferae* were irradiated or not with 200 $J/m^2$. Two hours after recovery in water ribosome were separated using a sucrose gradient as described in Fig. 1 and under Methods. RNA was purified from fractions 3-12 that represent disomes-light polysomes as described in Methods. Next, mRNA was further purified using poly dT and sent to sequencing. After sequencing, reads mapped to ribosomal RNA units were removed from the analysis as described in Methods. To complete the data set, total mRNA was purified from the same cultures without further ribosome separation. Genes that are enriched in the fractions 3-12 with and without UV were identified first. Next, genes that are enriched only in irradiated fractions 3-12 were identified and analyzed using STRING to reveal modules that are likely to by induced by UV at the translation level.

### Total early UV polysome UV

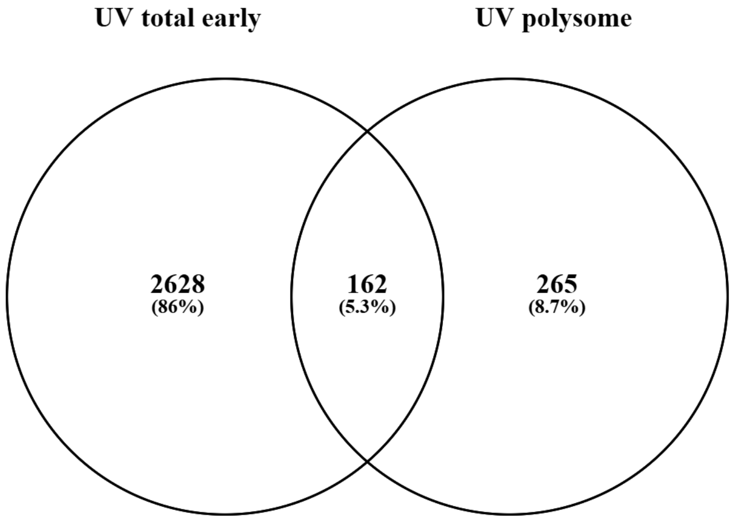


### Total early UV polysome control

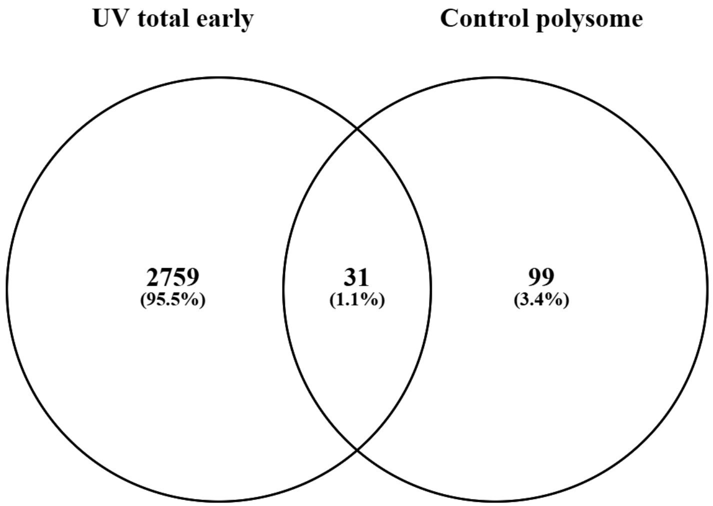

**Figure S9. Early UV-induced mRNA species are more associated with polysomes of UV-irradiated filaments than non-irradiated filaments.** Venn diagram created by Venny 2.1 of the genes that were induced by UV between 0-60 minutes of recovery time and mRNAs that are associated with polysomes 120 minutes after irradiation (left) or after 120 minutes of recovery with no irradiation (right). UV-induced genes tend to be more associated with polysomes after UV; Fisher's exact test: 0.0023. For early time points, total RNA was analyzed as described in the legend of Fig. S1. For polysome-associated mRNA, the experiment was done as described in the legend of Fig. 4.

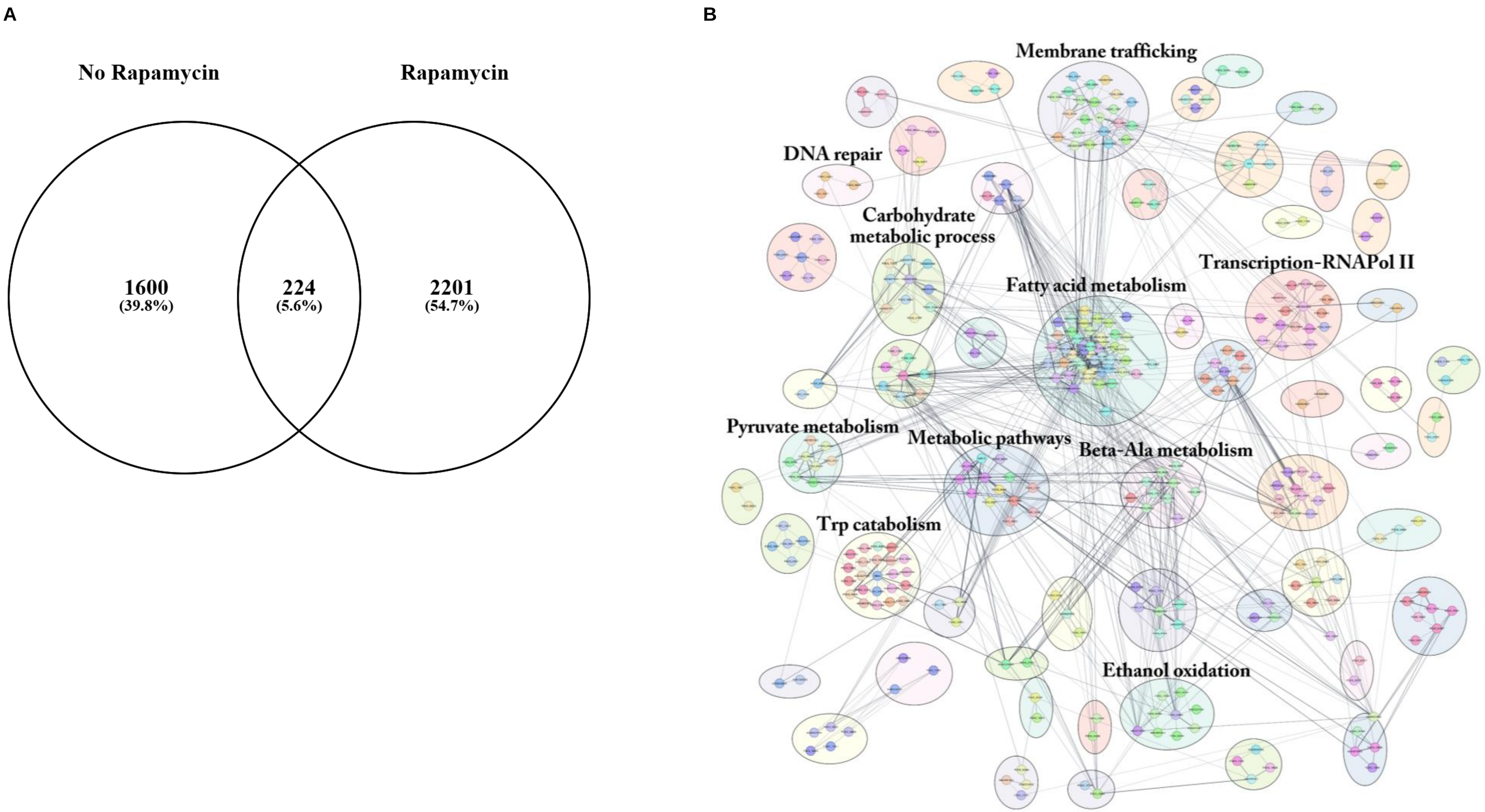
A
No Rapamycin
Rapamycin
1600
(39.8%)
224
(5.6%)
2201
(54.7%)
B
Membrane trafficking
DNA repair
Carbohydrate metabolic process
Transcription-RNAPol II
Fatty acid metabolism
Pyruvate metabolism
Metabolic pathways
Beta-Ala metabolism
Trp catabolism
Ethanol oxidation

**Figure S10. UV induction of ribosome biogenesis genes is severely affected by rapamycin**

(A). Venn diagram of *F. mangiferae* UV-induced genes with and without 5 μg/ml rapamycin. Conidia were inoculated into PDB for 13 h, treated with rapamycin for 1 h, irradiated with 200 J/m$^2$, and recovered for 30 minutes in water in the presence of rapamycin. RNA was purified from treated and untreated filaments. Induced genes under these conditions were compared with those induced by UV in the absence of rapamycin, as described in Milo et al.; Fungal Biology, 2024. (B). The genes up-regulated in the comparison of UV+rapamycin versus untreated were compared with those induced by rapamycin alone using Venn. F. oxysporum orthologs of the UV-specific induced genes were analyzed using STRING. Functional clusters presented using Cytoscape.

A

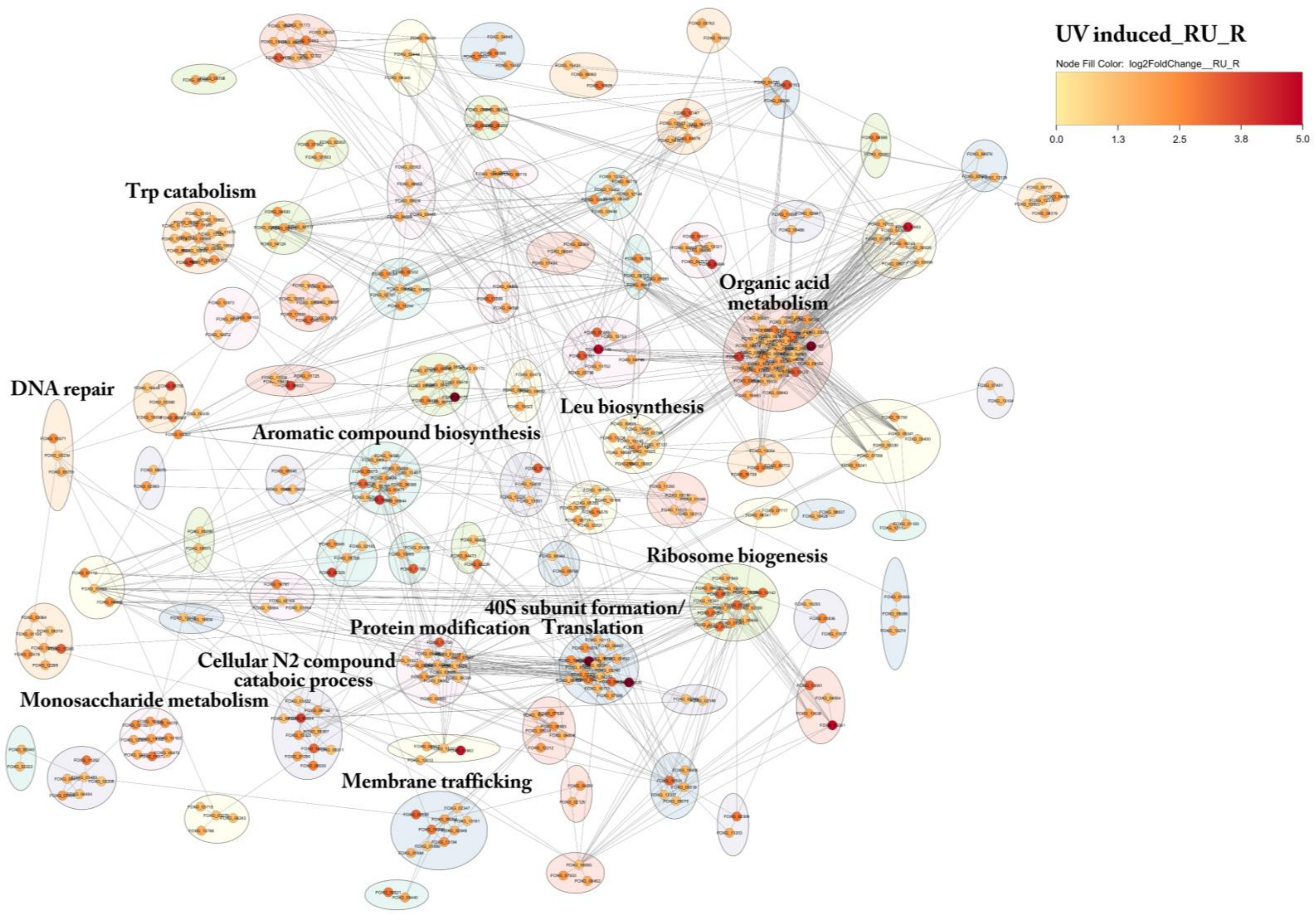
UV induced_RU_R
Node Fill Color: log2FoldChange__RU_R
0.0
1.3
2.5
3.8
5.0
Trp catabolism
DNA repair
Organic acid metabolism
Leu biosynthesis
Aromatic compound biosynthesis
Ribosome biogenesis
40S subunit formation/ Translation
Protein modification
Cellular N2 compound cataboic process
Monosaccharide metabolism
Membrane trafficking

B

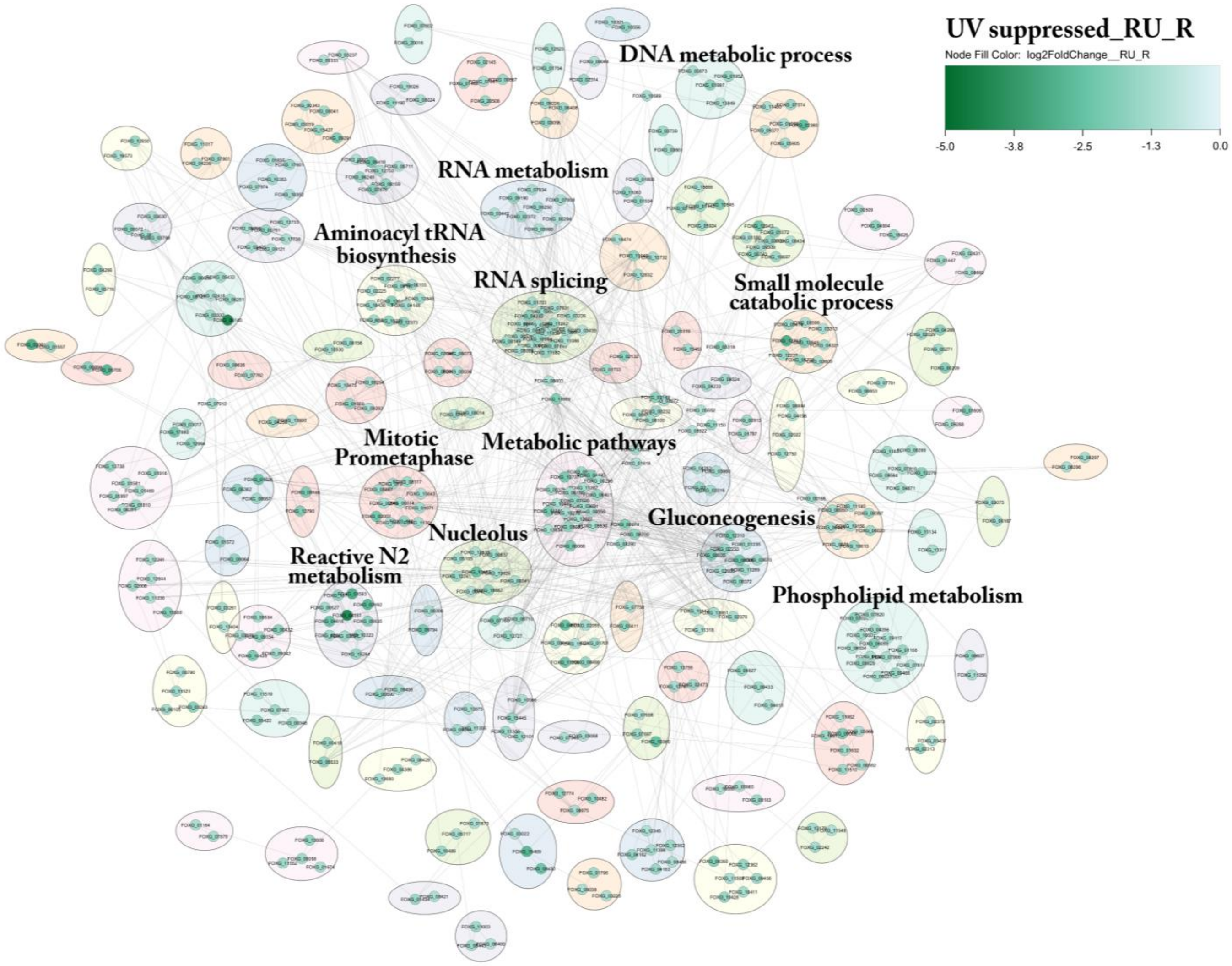

UV suppressed_RU_R
Node Fill Color: log2FoldChange__RU_R
-5.0
-3.8
-2.5
-1.3
0.0
DNA metabolic process
RNA metabolism
Aminoacyl tRNA biosynthesis
RNA splicing
Small molecule catabolic process
Mitotic Prometaphase
Metabolic pathways
Gluconeogenesis
Nucleolus
Reactive N2 metabolism
Phospholipid metabolism

**Figure S11. UV-differentially expressed genes under rapamycin exposure show a modest induction of translation-related genes.**

*F. oxysporum* orthologs of the *F. mangiferae* UV-induced (A) and reduced (B) genes by UV after pretreatment with rapamycin. Genes were clustered into functional groups using STRING. Functional clusters were presented using Cytoscape. The experiment was performed as described in the legend of Fig. S10.

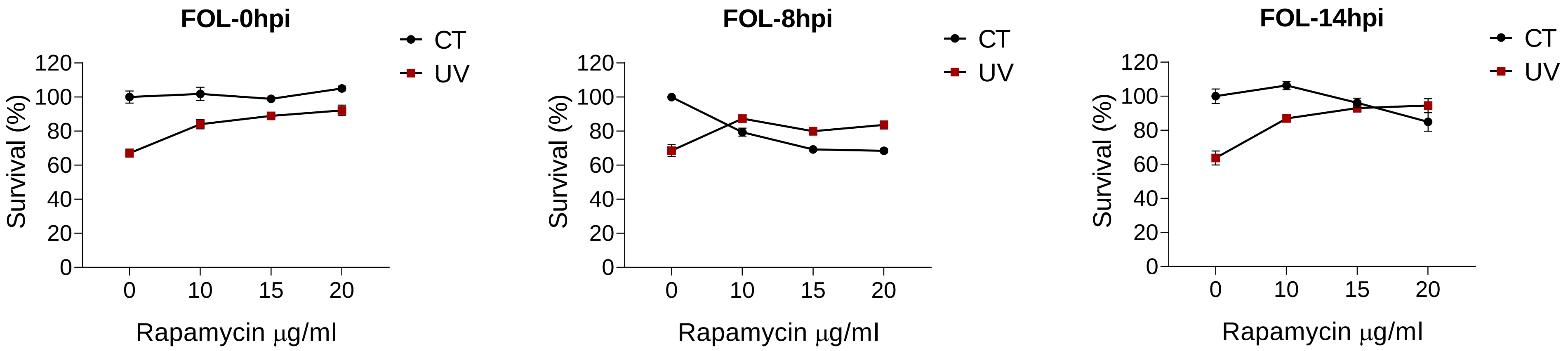


**Figure S12. UV rescues *F. oxysporum* from rapamycin toxicity.**

(A). *F. oxysporum* f.sp *lycopersici* (FOL) was inoculated at 0, 8, and 14 hours in PDB, then treated with rapamycin and UV as described in the legend of Fig. 5. Survival was calculated by dividing the number of colonies that grew after each treatment by the number of colonies in the culture that was not irradiated or treated with rapamycin (considered as 100%). Results are the average of 3 biological replicates, with error bars
